# Supra-physiological levels of Gibberellins/DELLAs alter the patterning, morphology and abundance of root hairs in root tips of *A. thaliana* seedlings

**DOI:** 10.1101/2021.07.15.452505

**Authors:** Iva McCarthy-Suárez

## Abstract

In spite of the known role of gibberellins (GAs), and of their antagonistic proteins, the DELLAs, in leaf hair production, no investigations, however, have assessed their hypothetical function in the production of root hairs. To this aim, the effects of supra-physiological levels of GAs/DELLAs on the spatial patterning of gene expression of the root hair (CPC) and root non-hair (GL2, EGL3 and WER) epidermal cell fate markers, as well as on the distribution, morphology and abundance of root hairs, were studied in root tips of 5-day-old *A. thaliana* seedlings. Results showed that excessive levels of GAs/DELLAs impaired the spatial patterning of gene expression of the root hair/non-hair epidermal cell fate markers, as well as the arrangement, shape and frequency of root hairs, giving rise to ectopic hairs and ectopic non-hairs, two-haired cells, two-tipped hairs, branched hairs, longer and denser hairs near the root tip under excessive DELLAs, and shorter and scarcer hairs near the root tip under excessive GAs. However, when the gai-1 (GA-insensitive-1) DELLA mutant protein was specifically over-expressed at the root epidermis, no changes in the patterning or abundance of root hairs occurred. Thus, these results suggest that, in seedlings of *A. thaliana,* the GAs/DELLAs might have a role in regulating the patterning, morphology and abundance of root hairs by acting from the sub-epidermal tissues of the root.

## 1. INTRODUCTION

The epidermal cell organization in roots of *A. thaliana* seedlings, consisting of single rows of hair-bearing cells (trichoblasts, which lay over the cleft between two cortical cells) alternating with double rows of hairless cells (atrichoblasts, which lay over just one cortical cell) has been shown to be genetically determined by a complex network of transcription factors and positional signals, such as CAPRICE (CPC), GLABRA2 (GL2), WEREWOLF (WER) and ENHANCER OF GLABRA3 (EGL3), and regulated by auxin, ethylene (ET), abscisic acid (ABA), nitric oxide (NO), brassinosteroids (BRs), cytokinins (CKs) and strigolactones (SLs)) (Silverman *et al.*, 1998; Cao *et al.,* 1999; Van Hengel *et al.,* 2004; Lombardo *et al.,* 2006; Kappusamy *et al.,* 2009; Schiefelbein *et al.,* 2009; Niu *et al.,* 2011; Salazar-Henao *et al.,* 2016). These hormones, in turn, seem to act downstream of the *GL2* gene network, permitting root cells to have fate plasticity, i.e., ability to change to the alternative differentiation route at a relatively late state, as it is not cell lineage, but position, and sometimes even a position-independent mechanism, what seems to continuously determine cell fate (Grierson and Schiefelbein, 2002; Schiefelbein *et al.,* 2009; Yu *et al.,* 2017). Moreover, these hormones mediate the changes in the root hair patterning associated to the plant responses to soil stress without altering the expression of the epidermal cell fate markers WER and GL2 (Schmidt *et al.,* 2000; Yang *et al.,* 2007; Martín-Rejano *et al.,* 2011).

Given that the GAs/DELLAs have a role in trichome (leaf hair) production in *A. thaliana* (Chien and Sussex, 1996; Traw and Bergelson, 2003) and participate in microtubule (MT) cytoskeleton organization (Locascio *et al.,* 2013), which is essential for the growth of trichomes and root hairs and for establishing the identity and shape of root cells (Bao *et al.,* 2001), and because there are no reports concerning the hypothetical implication of GAs/DELLAs in the root hair patterning, this study aimed to investigate the effect of excessive levels of these hormones on the distribution and abundance of root hairs in seedlings of *A*. *thaliana*. In addition, because changes in the levels of auxin, ET, ABA, NO, BRs and SLs have been correlated to alterations in root hair morphology in response to nutritional stresses such as low availability of P, B or Fe in the soil (longer and branched root hairs) (Schmidt *et al.,* 2000; Yang *et al.,* 2007; Martín-Rejano *et al.,* 2011), this work also aimed to determine whether the GAs / DELLAs might have a role in regulating the morphology of root hairs in seedlings of *A. thaliana*. To this aim, the spatial expression of the GUS or GFP-fused transcripts of the root hair (CPC) and non-hair (GL2, EGL3, WER) epidermal cell fate markers, as well as the arrangement, shape and density of root hairs, were studied in *A*. *thaliana* seedlings grown for 5 days under (or harbouring) excessive levels of GAs/DELLAs. Finally, to locate the tissue from which these hormones might hypothetically affect the patterning of root hairs, the root hair distribution was studied in 5-day-old mutant seedlings resulting from expressing the *gai-1* DELLA dominant allele in different tissues of the root (UAS (GAL4-UPSTREAM ACTIVATION SEQUENCE) expression directed system lines; Dr. Jim Haselhoff’s laboratory). Results of this study suggested that the GAs/DELLAs might be involved in regulating the patterning, morphology and abundance of root hairs in *A. thaliana* seedlings.

## 2. MATERIALS and METHODS

### 2.1. Plant Material and Growth Conditions

*Arabidopsis thaliana* Col (0) seeds were sterilized (70 % Ethanol (v/v) and 0.01 % Triton X-100 (v/v)), sown on half-strength MS medium plates (0.8 % (w/v) agar and 1 % (w/v) sucrose), stratified for 3-4 days (4°C, darkness), germinated, and grown vertically (22 °C; 5-7 days) under continuous white light (Percival growth chamber E-30B) (http://www.percival-scientific.com)as described by Lee and Schiefelbein (1999).

### 2.2. Hormone and Chemical treatments

Stock solutions of paclobutrazol (PAC, 10 mM in acetone 100 % (v/v)), GA_4_ (1 mM in 100 % ethanol (v/v)) or GA_3_ (50 mM in 100% ethanol (v/v)) were conveniently diluted and added to MS agar medium or water (in the case of liquid incubation experiments) to obtain a final concentration of 0.5 μM PAC, 1 μM GA_4_, and 30 μM GA_3_.

### 2.3. Mutant Lines

The spatial patterning of gene expression of the hair (CPC) and non-hair (GL2, EGL3, WER) epidermal cell fate markers in roots of *A. thaliana* seedlings was studied by using their GUS or GFP-fused promoter lines (*CPCpro::GUS*, *GL2pro::GUS*, *EGL3pro::GUS* and *WERpro::GFP*) as well as those derived from crossing lines harbouring constitutively excessive levels of GAs / DELLAs with the *GL2pro::GUS* line (*GID1b ox* x *GL2pro::GUS*, *gai-1* x *GL2pro::GUS*, *HSp::gai-1* x *GL2pro::GUS*, *pGAI::gai-1:GR* x *GL2pro::GUS* and *SCR::gai-1:GR* x *GL2pro::GUS* (L*er* x *GL2pro::GUS* background)). The effect of transient increases in the levels of the gai-1 dominant DELLA on the root hair distribution in *A. thaliana* seedlings was examined by using the heat-shock inducible *HSp::gai-1* (which over-expresses the DELLA gai-1 upon heat shock) and dexametasone (DEXA)-inducible *pGAI::gai-1*:GR and *SCR::gai-1*:GR (with glucocorticoid-binding domain) mutant lines. The *HSp::gai-1* mutant seedlings were grown at 37°C for 4 h (heat shock) and then at 22°C for 2 h (recovery period), whereas the *pGAI::gai-1:GR* and *SCR::gai-1:GR* mutant seedlings were incubated in 0.1, 0.2, 0.5, 1.2 or 10 μM DEXA for a minimum of 6h. The root hair distribution was also studied in mutants with excessive levels of GAs/DELLAs (*gai-1*, *GAI-ox* (GAI-over-expressing), *QD (quadruple DELLA mutant)*, *5X (quintuple DELLA mutant)*, GID1b-*ox* (which over-expresses the GA receptor GID1b (GIBBERELLIN INSENSITIVE DWARF1), in mutants over-expressing gai-1 in different tissues of the root [*ML1::gai-1* (epidermis) and UAS expression directed system (GAL4-UPSTREAM ACTIVATION SEQUENCE) mutants: *UAS::gai-1* x C24 (control, background); *UAS::gai-1* x J0951 (epidermis of the meristematic zone (MZ)); *UAS::gai-1* x J2812 (MZ epidermis and cortex); *UAS::gai-1* x N9142 (cortex of the elongation zone (EZ)); UAS::*gai-1* x M0018 (MZ cortex and endodermis); *UAS::gai-1* x J0571 (MZ cortex and endodermis); *UAS::gai-1* x Q2393 (all tissues but the endodermis); *UAS::gai-1* x Q2500 (MZ endodermis/pericycle); *UAS::gai-1* x J0121 (EZ pericycle); *UAS::gai-1* x J0631 (all tissues of the EZ); *UAS::gai-1* x J3281 (vessels)], and in the *wer*, *cpc* and *35S::CPC* (cauliflower mosaic virus 35S promoter) mutants.

### 2.4. GUS activity assay

GUS (β-glucuronidase) staining of the *GL2pro::GUS*, *CPCpro::GUS* and *EGL3pro::GUS* reporter lines was performed as described by Frigerio *et al.,* (2006), but using 8mM instead of 2 mM potassium ferro/ferricyanide and incubating the seedlings (15 min to 2h) in the reaction mixture at 4 °C instead of 37 °C.

### 2.5. Microscopy

The patterning of the hair/non-hair epidermal cell types in roots of *A. thaliana* seedlings was studied by staining the roots with 0.67 mg/ml propidium iodide, by observing the root tips under a Nickon Eclipse E6000 microscope, and by calculating the percentage of hairs/non-hairs at the trichoblast/atrichoblast positions (Dr. Benedicte Desvoyes’ method). The *GL2pro::GUS* expression patterning in cross sections of root tips was studied on ultra-thin cross sections of plastic resin-embedded roots as previously described at Dr. Schiefelbein Lab Protocols (http://www.mcdb.lsa.umich.edu/labs/schiefel/protocols.html). Seedlings were included in 1% agarose in 0.1M sodium phosphate buffer, pH 6.8, and stained for GUS activity. Root-containing blocks were then cut, fixed with 4% para-formaldehyde in PBS, dehydrated in ethanol series (15%, 30%, 50%, 75%, 95% and 100%, 1h each), kept in 100% ethanol overnight, incubated in Technovit ® 7100 infiltration solution for 2 days, inserted in gelatine capsules, and embedded for 9 days in Technovit ® 7100 plastic resin (Heraeus Kultzer, Germany). Ultramicrotome (Ultracut E, Reichert Jung, Germany) cross sections of resin-embedded roots were then stained with 0.06% (w/v) toluidine blue and observed under a Nikon Eclipse E600 microscope. The *WERpro::GFP* expression was visualized by using a Leica Confocal Microscope (excitation: 488nm; detection: 500-530nm band-path filter for GFP).

## 3. RESULTS

### 3.1. Excessive levels of GAs/DELLAs altered the root hair patterning in seedlings of *A. thaliana*

To assess whether the GAs/DELLAs might have a role in the root hair patterning of *A. thaliana* seedlings, the spatial gene expression of the root hair (CPC) and non-hair (GL2, EGL3, WER) epidermal cell fate markers was studied in seedlings of the *GL2pro::GUS*, *CPCpro::GUS*, *EGL3pro::GUS* and *WERpro::GFP* transgenic lines grown for 5 days under supra-physiological levels of GAs/DELLAs (Fig. 1A). Results showed that growth under excessive levels of GAs/DELLAs altered the normal patterning of gene expression the root hair/non-hair epidermal cell fate markers (Fig. 1A). This was confirmed by analysing the spatial expression of *GL2* in the *GID1b-ox* (which over-expresses the GA receptor GID*1b*), *gai-1*, *HSp::gai-1* (which over-expresses the DELLA gai-1 upon exposure to heat (37°C, 4h)), and DEXA-inducible *pGAI::gai-1:GR* and *SCR::gai-1:GR* mutants (Fig. 1A). Moreover, the alteration of the *GL2pro::GUS* expression pattern under excessive levels of GAs/DELLAs was corroborated in ultra-thin sections of resin-embedded roots (Fig. 1B).

**Fig. 1A.**
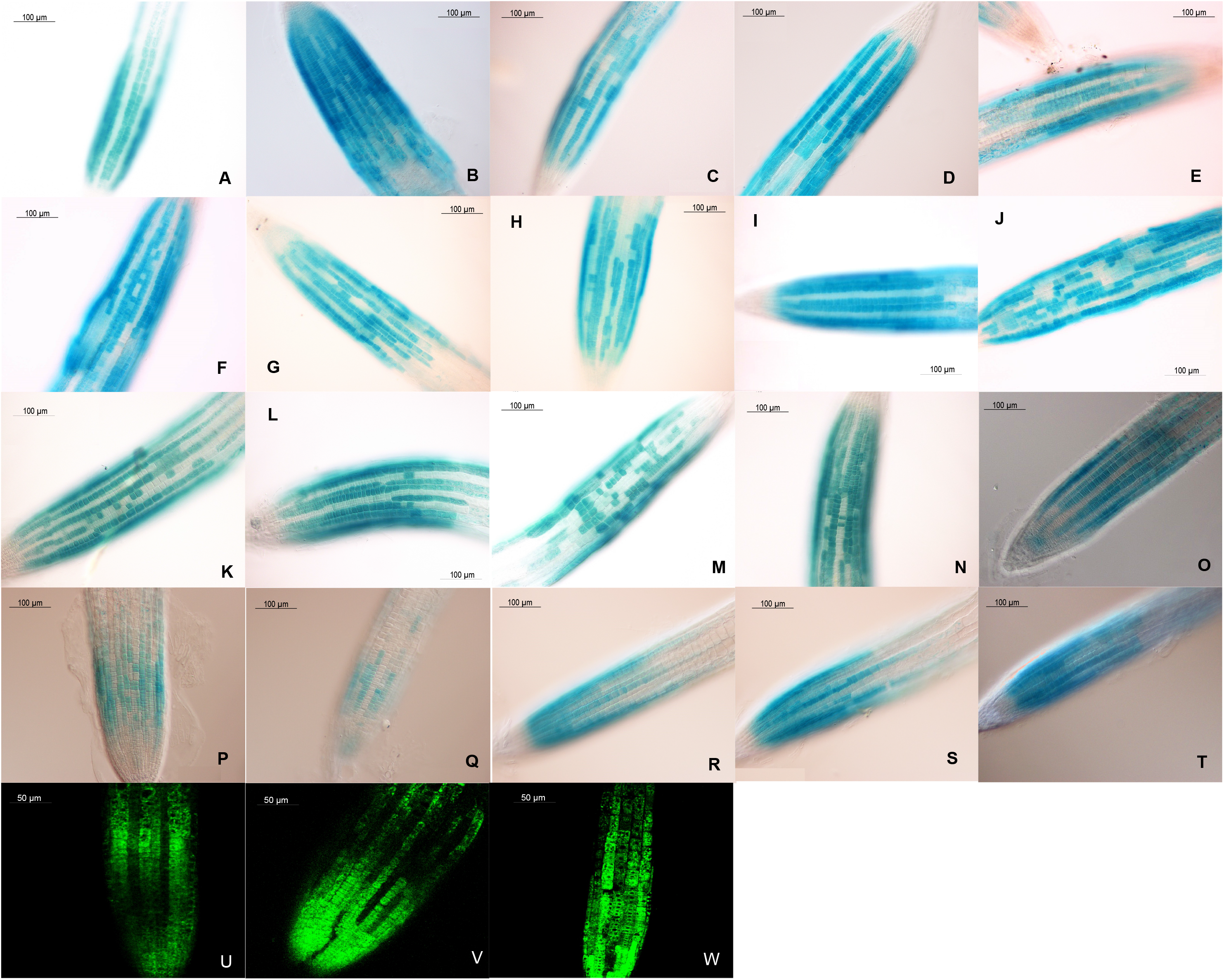
Spatial gene expression of the root hair (*CPC) and non-hair (GL2, *EGL3, WER) epidermal cell fate markers in 5-day-old A. thaliana seedlings grown under (or harbouring) excessive levels of GAs/DELLAs. **A)** *GL2pro::GUS* (MS), 20X; **B)** *GL2pro::GUS* (0.5 μM PAC), 20X; **C)** *GL2pro::GUS* (1 μM GA_4_), 20X; **D)** *GL2pro::GUS* (30 μM GA_3_), 20X; **E)** L*er* x *GL2pro::GUS* (MS), 20X; **F)** *gai-1* x *GL2pro::GUS* (MS), 20X; **G)** GID1b-*ox* x *GL2pro::GUS* (24h in H_2_O; liquid incubation experiment; leaky line), 20X; **H)** GID1b-*ox* x *GL2pro::GUS* (24h in 1 μM GA_4_; liquid incubation experiment), 20X; **I)** *HSp::gai-1* x *GL2pro::GUS* (22°C, 4h), 20X; **J)** *HSp::gai-1* x *GL2pro::GUS* (37°C, 4h), 20X; **K)** *pGAI::gai-1:GR* x *GL2pro::GUS* (24h in MS; leaky line), 20X; **L)** *pGAI::gai-1:GR* x *GL2pro::GUS* (24h in 10 μM DEXA), 20X; **M)** *SCR::gai-1:GR* x *GL2pro::GUS* (24h in MS; leaky line), 20X; **N)** *SCR::gai-1:GR* x *GL2pro::GUS* (24h in 10 μM DEXA), 20X; **O)** *CPCpro::GUS* (MS), 20X; **P)** *CPCpro::GUS* (0.5 μM PAC), 20X; **Q)** *CPCpro::GUS* (1 μM GA_4_), 20X; **R)** *EGL3pro::GUS* (MS), 20X; **S)** *EGL3pro::GUS* (0.5 μM PAC), 20X; **T)** *EGL3pro::GUS* (1 μM GA_4_), 20X; **U)** *WERpro::GFP* (MS), 40X; **V)** *WERpro::GFP* (0.5 μM PAC), 40X; **W)** *WERpro::GFP* (1 μM GA_4_), 40X. In control seedlings, *GL2* is expressed in root non-hair (atrichoblast) cells. *CPC protein is expressed in root non-hair cells, but migrates to root hair cells. *EGL3 protein is expressed in root hair cells, but migrates to root non-hair cells.

**Fig. 1B.**
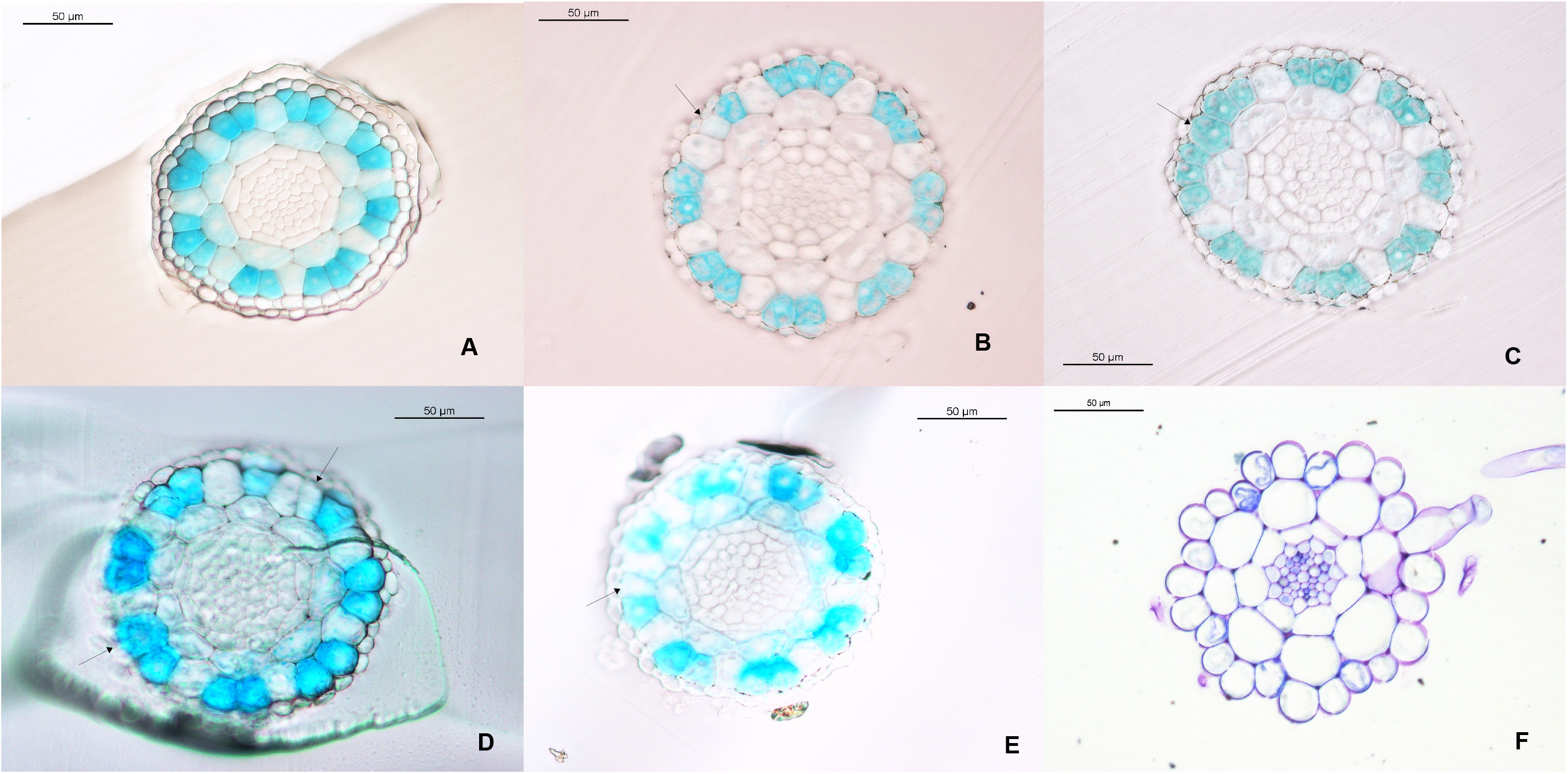
Spatial gene expression of the root non-hair epidermal cell fate marker GL2 in cross sections of resin-embedded roots of A. thaliana seedlings grown for 5 days under excessive levels of GAs/DELLAs. **A)** *GL2pro::GUS* (MS); **B)** *GL2pro::GUS* (0.5 μM PAC): Lack of *GL2* expression in an atrichoblast cell; **C)** *GL2pro::GUS* (0.5 μM PAC): Ectopic expression of *GL2* in a trichoblast cell; **D)** *GL2pro::GUS* (1 μM GA_4_): Ectopic expression of *GL2* in a trichoblast cell and lack of *GL2* expression in an atrichoblast cell; **E)** *GL2pro::GUS* (1 μM GA_4_): Lack of *GL2* expression in an atrichoblast cell; **F)** Ectopic root hair cell in *gai-1*. Magnification: 40X.

An analysis of the distribution of root hairs and non-hairs relative to their position over the cortex cells showed that excessive levels of GAs/DELLAs impaired the correct positioning of the root hairs/non-hairs, giving rise to ectopic root hairs (at the atrichoblast position) and ectopic root non-hairs (at the trichoblast position) (Figs. 2A and 2B; Tables 1 and 2). Interestingly, treatment with GA_4_ (1 μM) reduced the percentage of ectopic root hairs in the hairy *35S::CPC* mutant, whereas treatment with PAC (0.5 μM) slightly decreased the percentage of ectopic root non-hairs in the bald *cpc* mutant (Table 2). In accordance with these changes, growing *A. thaliana* seedlings under supra-physiological levels of GAs/DELLAs for 5 days altered the arrangement of root hairs in root tips, giving rise to ectopic hairs (in a non-hair row), ectopic non-hairs (in a hair row) and adjacent hair rows (Figs. 3A and 3B). This was confirmed in the *gai-1*, *QD*, *5X*, GID*1b-ox*, *pGAI::gai-1:GR* and *SCR::gai-1:GR* mutants (Figs. 3A and 3B).

**Fig. 2A.**
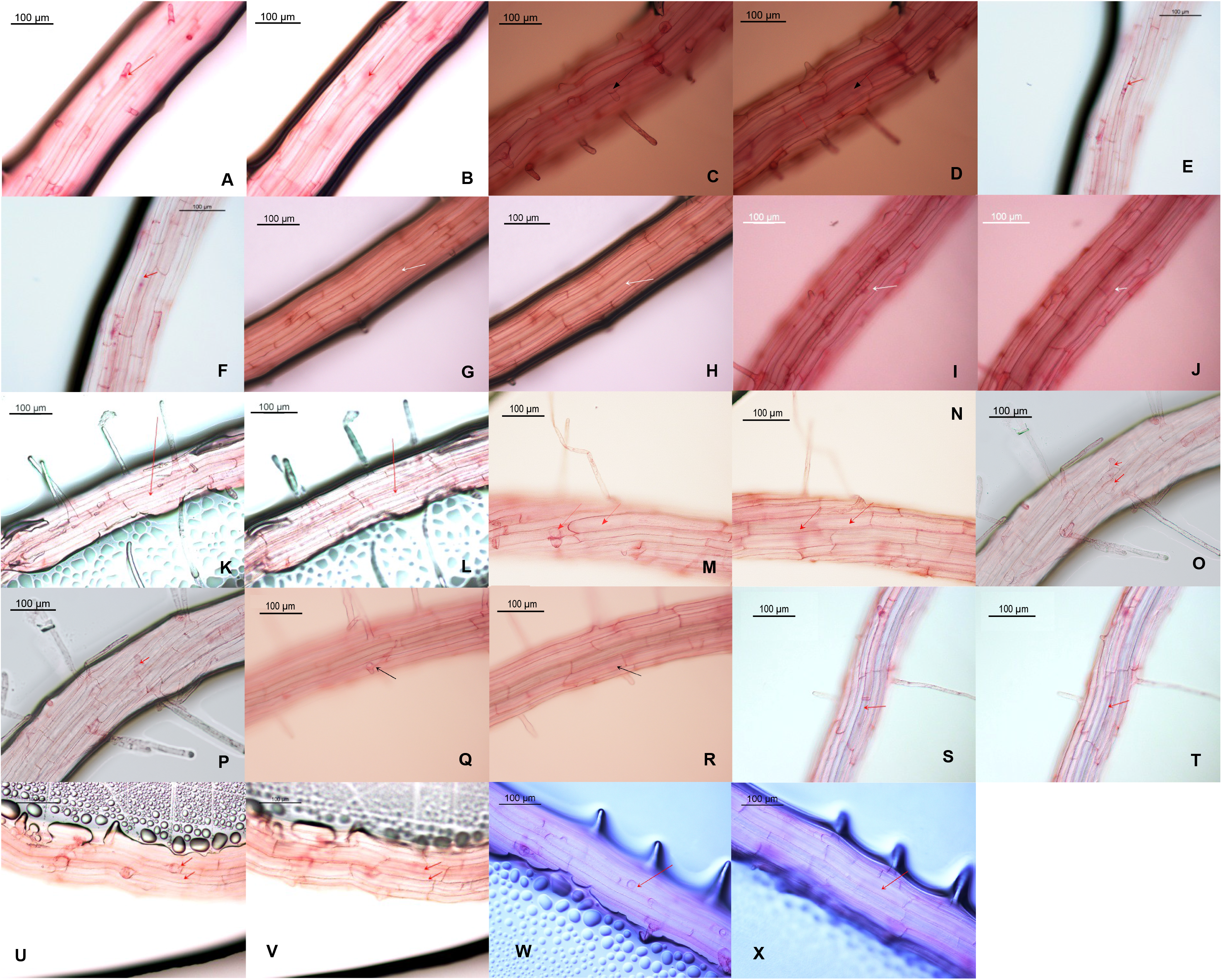
Ectopic root hairs and ectopic root non-hairs in root tips of 5-day-old A. thaliana seedlings grown under (or harbouring) excessive levels of GAs/DELLAs. **A)** Col(0) (MS) correct hair (epidermis); **B)** Col(0) (MS), correct hair (cortex); **C)** Col(0) (0.5 μM PAC) ectopic hair (epidermis); **D)** Col(0) (0.5 μM PAC) ectopic hair (cortex); **E)** Col(0) (1 μM GA_4_) ectopic hair (epidermis); **F)** Col(0) (1 μM GA_4_) ectopic hair (cortex); **G)** Col(0) (MS) correct non-hair (epidermis); **H)** Col(0) (MS) correct non-hair (cortex); **I)** Col(0) (0.5 μM PAC) ectopic non-hair (epidermis); **J)** Col(0) (0.5 μM PAC) ectopic non-hair (cortex); **K)** Col(0) (1 μM GA_4_) ectopic non-hair (epidermis); **L)** Col(0) (1 μM GA_4_) ectopic non-hair (cortex); **M)** L*er*, correct hair and non-hair (epidermis); **N)** L*er*, correct hair and non-hair (cortex); **O)** *gai-1*, ectopic hair (epidermis); **P)** *gai-1*, ectopic hair (cortex); **Q)** *QD,* ectopic hair (epidermis); **R)** *QD,* ectopic hair (cortex); **S)** *5X,* ectopic non-hair (epidermis); **T)** *5X,* ectopic non-hair (cortex); **U)** *pGAI::gai-1:GR* (10 μM DEXA), ectopic non-hair and ectopic hair (epidermis); **V)** *pGAI::gai-1:GR* (10 μM DEXA), ectopic non-hair and ectopic hair (cortex); **W)** *HSp::gai-1,* 2d after heat shock (37°C, 4h), ectopic hair (epidermis); **X)** *HSp::gai-1,* 2d after heat-shock (37°C, 4h), ectopic hair (cortex). Magnification: 20X.

**Fig. 2B.**
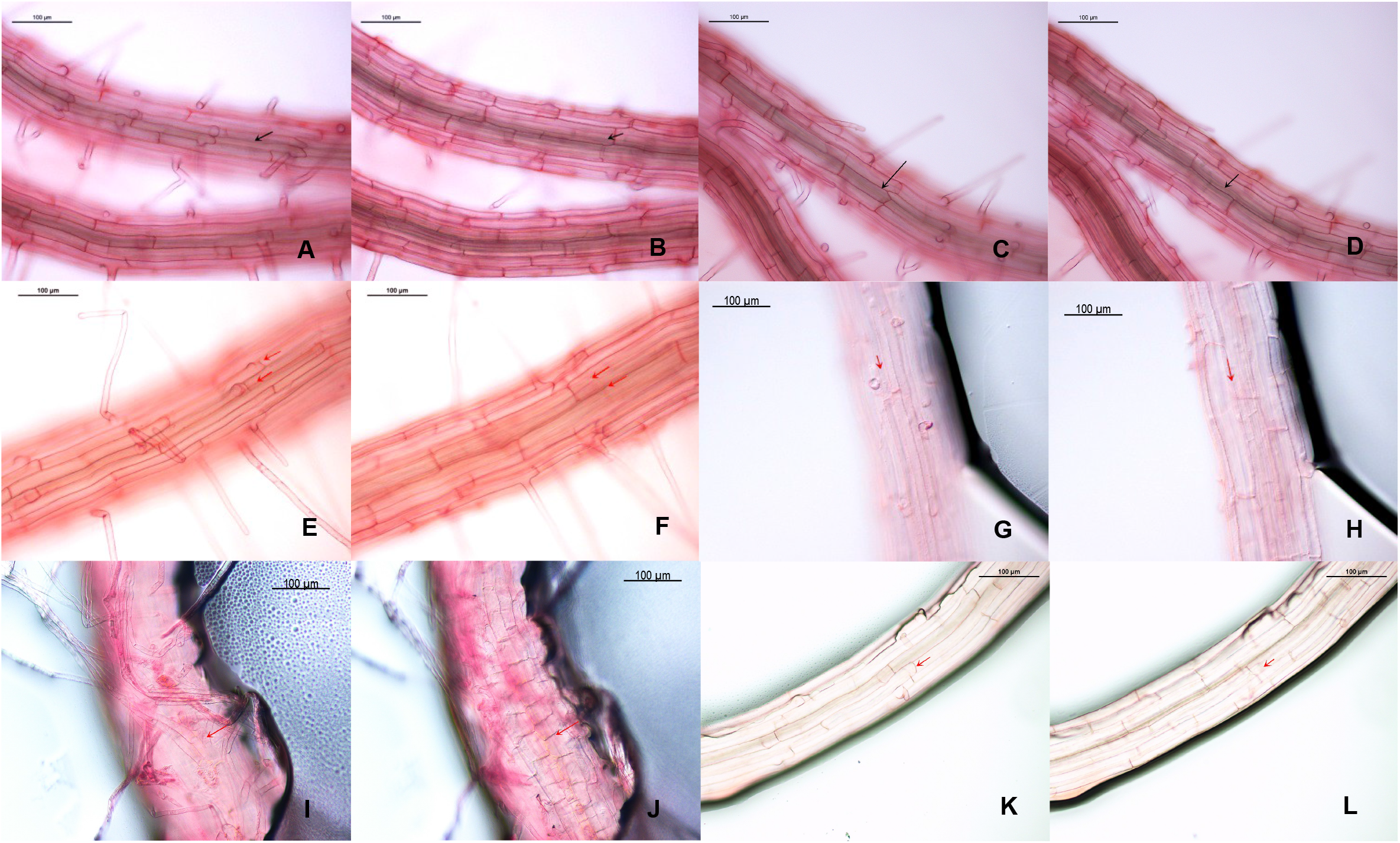
Ectopic hairs and non-hairs in root tips of 5-day-old A. thaliana seedlings over-expressing the DELLA gai-1 in different tissues of the root. **A)** *UAS::gai-1* x J2812, ectopic hairs (epidermis); **B)** *UAS::gai-1* x J2812, ectopic hairs (cortex); **C)** *UAS::gai-1* x J2812 (ectopic non-hair, epidermis); **D)** *UAS::gai-1* x J2812 (ectopic non-hair, cortex); **E)** *UAS::gai-1* x Q2393 (ectopic hair, epidermis); **F)** UAS::*gai-1* x Q2393 (ectopic hair, cortex); **G)** *UAS::gai-1* x Q2393 (ectopic non-hair, epidermis); **H)** *UAS::gai-1* x Q2393 (ectopic non-hair, cortex); **I)** *UAS::gai-1* x Q2500 (ectopic hair, epidermis); **J)** *UAS::gai-1* x Q2500 (ectopic hair, cortex); **K)** *UAS::gai-1* x J0121 (ectopic non-hair, epidermis); **L)** *UAS::gai-1* x J0121 (ectopic non-hair, cortex). Magnification: 20X.

**Table 1.** Distribution of hair and non-hair cells at the trichoblast/atrichoblast positions in roots of 5-day-old A. thaliana seedlings grown under (or harbouring) excessive levels of GAs/DELLAs. Numbers in parenthesis refer to the number of cells analysed. At least 15-20 roots were used per treatment.

|  | Trichoblast position |  | Atrichoblast position |  |
| --- | --- | --- | --- | --- |
|  | Hair cell<br>(%) | Non-hair cell<br>(%) | Hair cell<br>(%) | Non-hair cell<br>(%) |
| Col (0) (MS) | 97.5 ± 0.7 (73) | 2.5 ± 0.7 (2) | 0 ± 0 (0) | 100 ± 0 (75) |
| PAC (0.5 µM) | 77.4 ± 5.7 (109) | 22.6 ± 5.7 (32) | 36.7 ± 8.2 (80) | 63.3 ± 8.2 (138) |
| GA<sub>4</sub> (1 µM) | 81 ± 2.7 (83) | 19 ± 2.7 (20) | 12.5 ± 3.5 (5) | 87.5 ± 3.5 (35) |
| PAC (0.5 µM) + GA<sub>4</sub> (1 µM) | 94 ± 4.2 (71) | 6 ± 4.2 (5) | 5 ± 2.8 (4) | 95 ± 2.8 (71) |
| Ler | 95.8 ± 2.2 (167) | 4.2 ± 2.2 (7) | 4.5 ± 3.5 (6) | 95.5 ± 3.5 (120) |
| *gai-1* | 82.7 ± 4.5 (75) | 17.3 ± 4.5 (16) | 40.4 ± 5 (55) | 59.6 ± 5 (81) |
| QD | 78.8 ± 4.5 (126) | 21.2 ± 4.5 (34) | 24 ± 4.9 (38) | 76 ± 4.9 (122) |
| *pGAI::gai-1:GR* (MS) | 93.5 ± 2.1 (41) | 6.5 ± 2.1 (3) | 25 ± 7.1 (10) | 75 ± 7.1 (30) |
| *pGAI::gai-1:GR* (10 µM DEXA) | 78 ± 2.8 (38) | 22 ± 2.8 (11) | 50.5 ± 6.4 (22) | 49.5 ± 6.4 (22) |
| *SCR::gai-1:GR* (MS) | 83.8 ± 3.3 (36) | 16.2 ± 3.3 (7) | 35 ± 6.2 (13) | 65 ± 6.2 (25) |
| *SCR::gai-1:GR* (0.1 µM DEXA) | 67 ± 12.7 (30) | 33 ± 12.7 (15) | 15 ± 7.1 (6) | 85 ± 7.1 (34) |

**Table 2.** Percentage of ectopic root hair/non-hair cells at the trichoblast/atrichoblast positions in roots of 5-day-old A. thaliana seedlings grown under (or harbouring) excessive levels of GAs/DELLAs. *Seedlings analysed at 48h after a heat-shock experiment (37°C, 4h). *GL2pro::GUS* (22°C): control seedlings grown at 22°C for 4h. *GL2pro::GUS* (37°C): control seedlings grown at 37°C for 4h (ectopic root hair cells might have appeared due to heat stress). *HSp::gai-1* x *GL2pro::GUS* (37°C): inducible *gai-1* mutant seedlings grown at 37°C for 4h. The number of ectopic root hair and non-hair cells from a single experiment is shown in parenthesis.

|  | Trichoblast position |  | Atrichoblast position |  |
| --- | --- | --- | --- | --- |
|  | Number of epidermal cells examined | % Ectopic root non-hair cells | Number of epidermal cells examined | % Ectopic root hair cells |
| *Ler* | 41 | 2 (1) | 30 | 7 (2) |
| *5X* | 20 | 35 (7) | 20 | 0 (0) |
| *GAI-ox* | 15 | 7 (1) | 15 | 53 (8) |
| *GL2pro::GUS* (22°C) * | 30 | 0 (0) | 29 | 0 (0) |
| *GL2pro::GUS* (37°C)* | 30 | 7 (2) | 30 | 30 (9) |
| *Hsp::gai-1* x *GL2pro::GUS* (37°C)* | 29 | 28 (8) | 27 | 44 (12) |
| *wer* | 28 | 32 (9) | 36 | 50 (18) |
| *wer* (1 µM GA<sub>4</sub>) | 30 | 20 (6) | 29 | 72 (21) |
| *cpc* | 28 | 64 (18) | 28 | 21 (6) |
| *cpc* (0.5 µM PAC) | 28 | 54 (15) | 28 | 4 (1) |
| *35S::CPC* | 30 | 23 (7) | 30 | 60 (18) |
| *35S::CPC* (1 µM GA<sub>4</sub>) | 30 | 13 (4) | 29 | 17 (5) |
| *SCR::gai-1:GR* (MS) | 21 | 19 (4) | 20 | 30 (6) |
| *SCR::gai-1:GR* (0.2 µM DEXA) | 19 | 21 (4) | 19 | 26 (5) |
| *SCR::gai-1:GR* (0.5 µM DEXA) | 20 | 30 (6) | 18 | 22 (4) |
| *SCR::gai-1:GR* (1.2 µM DEXA) | 10 | 20 (2) | 10 | 40 (4) |
| *UAS::gai 1* x *C24* (control) | 50 | 0 (0) | 44 | 0 (0) |
| *ML1::gai-1* | 41 | 2 (1) | 30 | 3 (1) |
| *UAS::gai-1* x *J0951* | 60 | 0 (0) | 60 | 0 (0) |
| *UAS::gai-1* x *J2812* | 30 | 10 (3) | 30 | 50 (15) |
| *UAS::gai-1* x *Q2393* | 9 | 22 (2) | 16 | 63 (10) |

**Fig. 3A.**
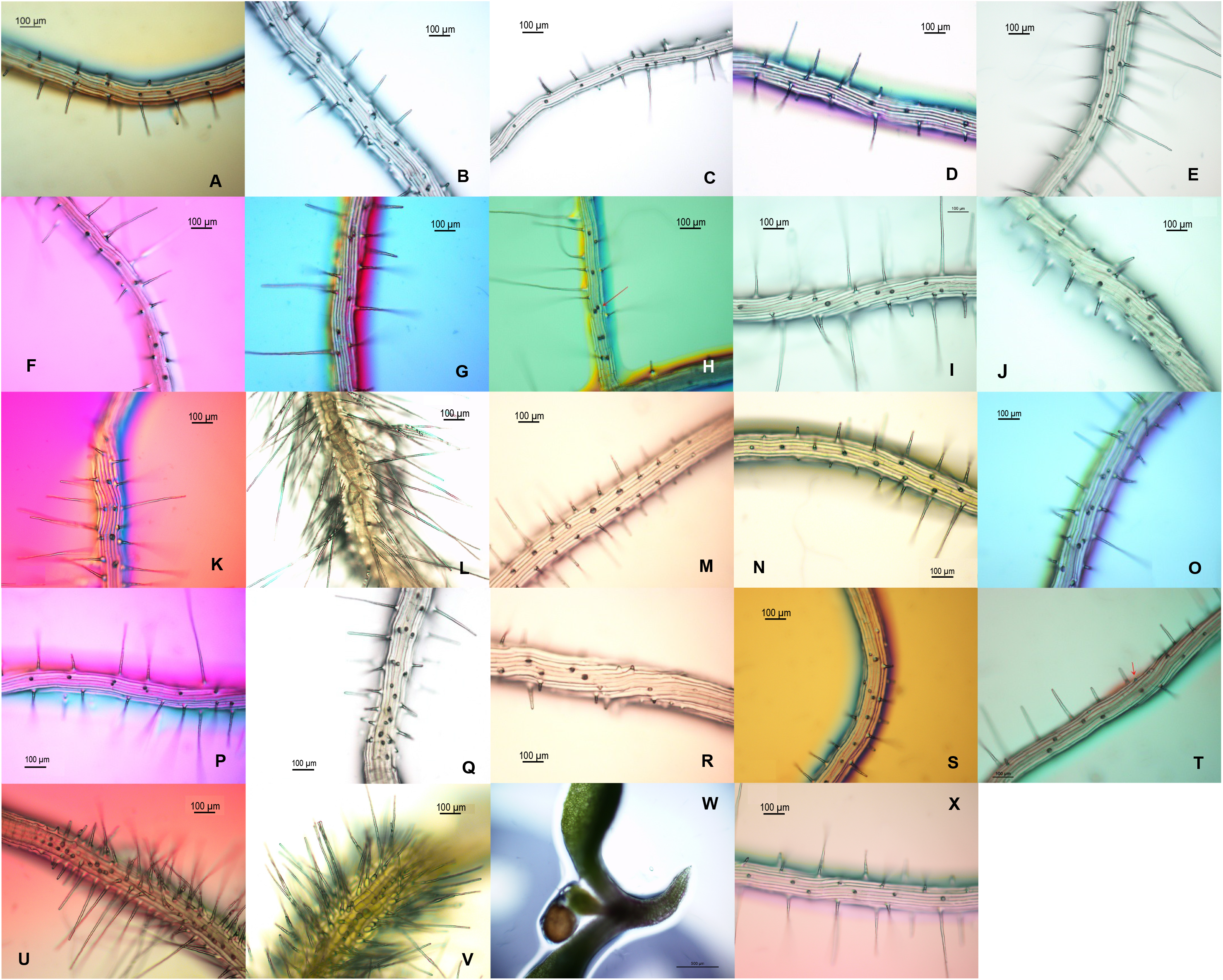
Arrangement of hairs in root tips of 5-day-old A. thaliana seedlings grown under (or harbouring) excessive levels of GAs/DELLAs. **A)** Col(0) (MS), 10X; **B)** Col(0) (0.5 μM PAC), 10X; **C)** Col(0) (30 μM GA_3_), 10X; **D)** L*er*, 10X; **E)** *gai-1,* 10X; **F)** *QD*, 10X; **G)** *5X,* 10X; **H)** *GID1b*-*ox* (MS, leaky line), lateral root, 10X; **I)** *pGAI::gai-1:GR* (30h in MS; leaky line), 10X; **J)** *pGAI::gai-1:GR* (30h in 10 μM DEXA), 10X; **K)** *SCR::gai-1:GR* (72h in MS; leaky line), 10X; **L)** *SCR::gai-1:GR* (48h in 10 μM DEXA), 10X; **M)** *UAS::gai-1* x C24, 10X; **N)** *UAS::gai-1* x J0951, 10X; **O)** *UAS::gai-1* x J2812, 10X; **P)** *UAS::gai-1* x N9142, 10X; **Q)** *UAS::gai-1* x M0018, 10X; **R)** *UAS::gai-1* x Q2500, 10X; **S)** *UAS::gai-1* x Q2393, 10X; **T)** *UAS::gai-1* x J0121, 10X; **U)** *UAS::gai-1* x J0631, 10X; **V)** *UAS::gai-1* x J0571, 10X; **W)** *UAS::gai-1* x J3281, 4X; **X)** *ML1::gai-1*, 10X.

**Fig. 3B.**
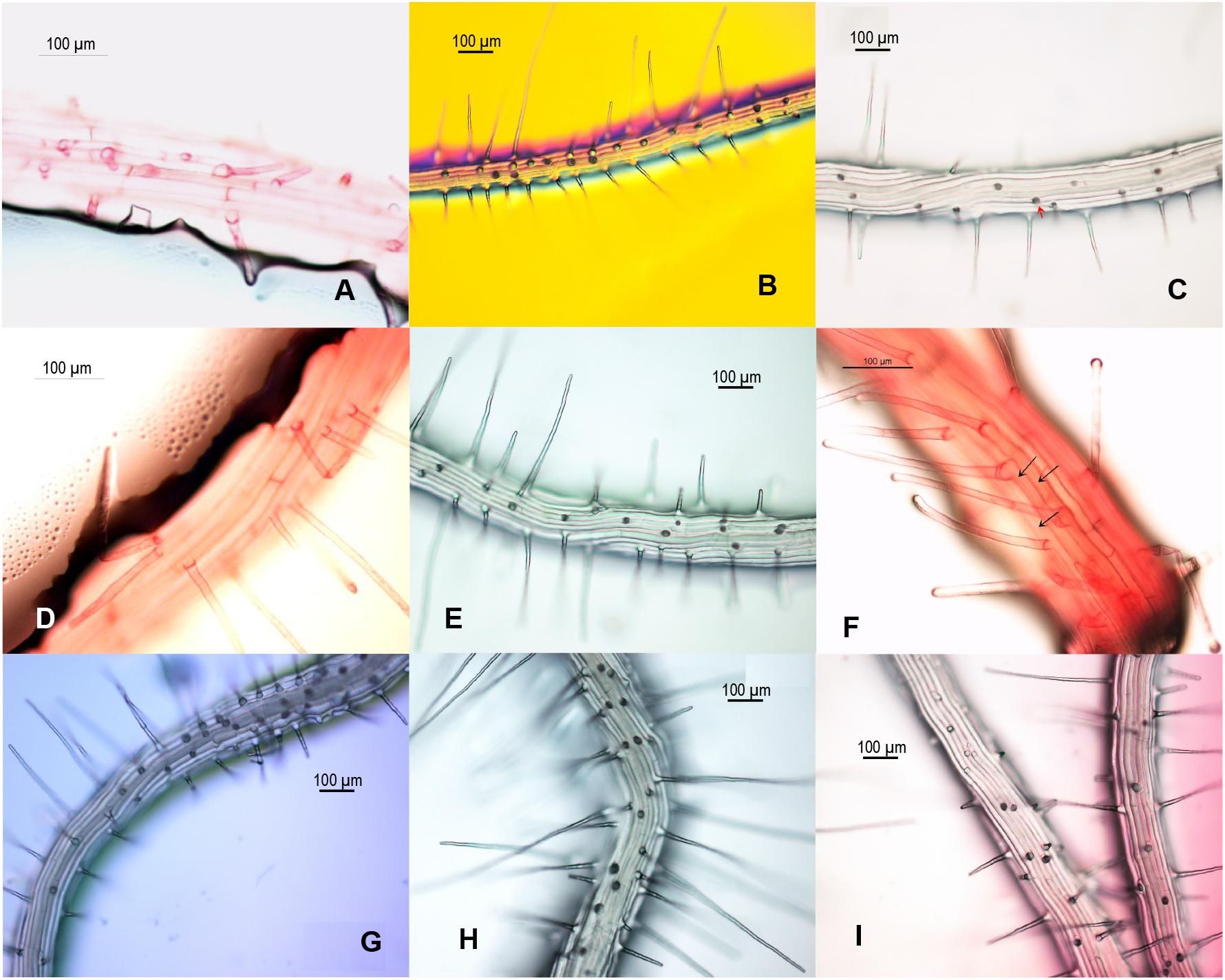
Adjacent hair rows in root tips of 5-day-old A. thaliana seedlings grown under (or harbouring) excessive levels of GAs/DELLAs. **A)** Col(0) (0.5 μM PAC), 20X; **B)** *gai-1* (lateral root), 10X; **C)** *QD,* 10X; *HSp::gai-1,* 2 days after heat shock (37°C, 4h), 20X; **E)** *pGAI::gai-1:GR* (MS, leaky line), 10X; **F)** *SCR::gai-1:GR* (MS, leaky line), 20X; **G)** *UAS::gai-1* x J2812, 10X; **H)** *UAS::gai-1* x M0018, 10X; **I)** *UAS::gai-1* x Q2393, 10X.

To ascertain from which particular tissue of the root the GAs/DELLAs might be affecting the root hair patterning, the distribution of root hairs and the positioning of the root hair/non-hair cells over the root cortex cells were studied in *A. thaliana* transgenic seedlings expressing the *gai-1* DELLA allele in different tissues of the root (Figs. 2B, 3A and 3B; Table 2). Results showed that the root hair distribution changed when *gai-1* was over-expressed at the cortex, endodermis or pericycle of the meristematic (MZ) or elongation (EZ) zones of the root (J2812 ≫ *gai-1*, M0018 ≫ *gai-1*, Q2500 ≫ *gai-1*, J0121 ≫ *gai-1*, Q2393 ≫ *gai-1* and J0631 ≫ *gai-1* lines), but not when *gai-1* was over-expressed at the root epidermis (J0951 ≫ *gai-1* and *ML1::gai-1* lines) (Fig. 3A). Moreover, ectopic hairs, ectopic non-hairs and adjacent hair rows appeared when *gai-1* was over-expressed at the cortex (J2812 ≫ *gai-1* and N9142 ≫ *gai-1* lines), endodermis (M0018 ≫ *gai-1* and J0571 ≫ *gai-1* lines) or pericycle (J0121 ≫ *gai-1* line) of the root, or in all root tissues but the endodermis (Q2393 ≫ *gai-1* line) (Figs. 2B, 3A and 3B). However, when *gai-1* was over-expressed at the root vessels (J3281 ≫ *gai-1* line), the production of root hairs and the growth of the root stopped (Fig. 3A).

### 3.2. Excessive levels of GAs/DELLAs altered the morphology, length and abundance of hairs in root tips of *A. thaliana* seedlings

Excessive levels of GAs/DELLAs also modified the morphology of trichoblasts and root hairs in root tips of *A. thaliana* seedlings, frequently giving rise to two-haired cells, two-tipped hairs and branched hairs (Fig 4). In addition, excessive levels of GAs/DELLAs altered the length and density of root hairs. Whereas high levels of DELLAs increased the length and number of hairs near the root tip, high levels of GAs had the opposite effect (Figs. 5A and 5B; Table 3). Moreover, hair abundance in root tips of *A. thaliana* seedlings increased when gai-1 was over-expressed at the cortex (J2812 ≫ *gai-1*), endodermis (M0018 ≫ *gai-1*) or pericycle (J0121 ≫ *gai-1*) of the root, but not when gai-1 was over-expressed at the epidermis of the root MZ (J0951 ≫ *gai-1*) or the cortex of the root EZ (N9142 ≫ *gai-1*) (Table 3). Also, treatment of the bald *cpc* mutant with PAC slightly increased the hair frequency (and length) near the root tip, whereas treatment of the hairy *wer* and *35S::CPC* mutants with GA_4_ reduced it (Fig. 5B; Table 3).

**Fig. 4.**
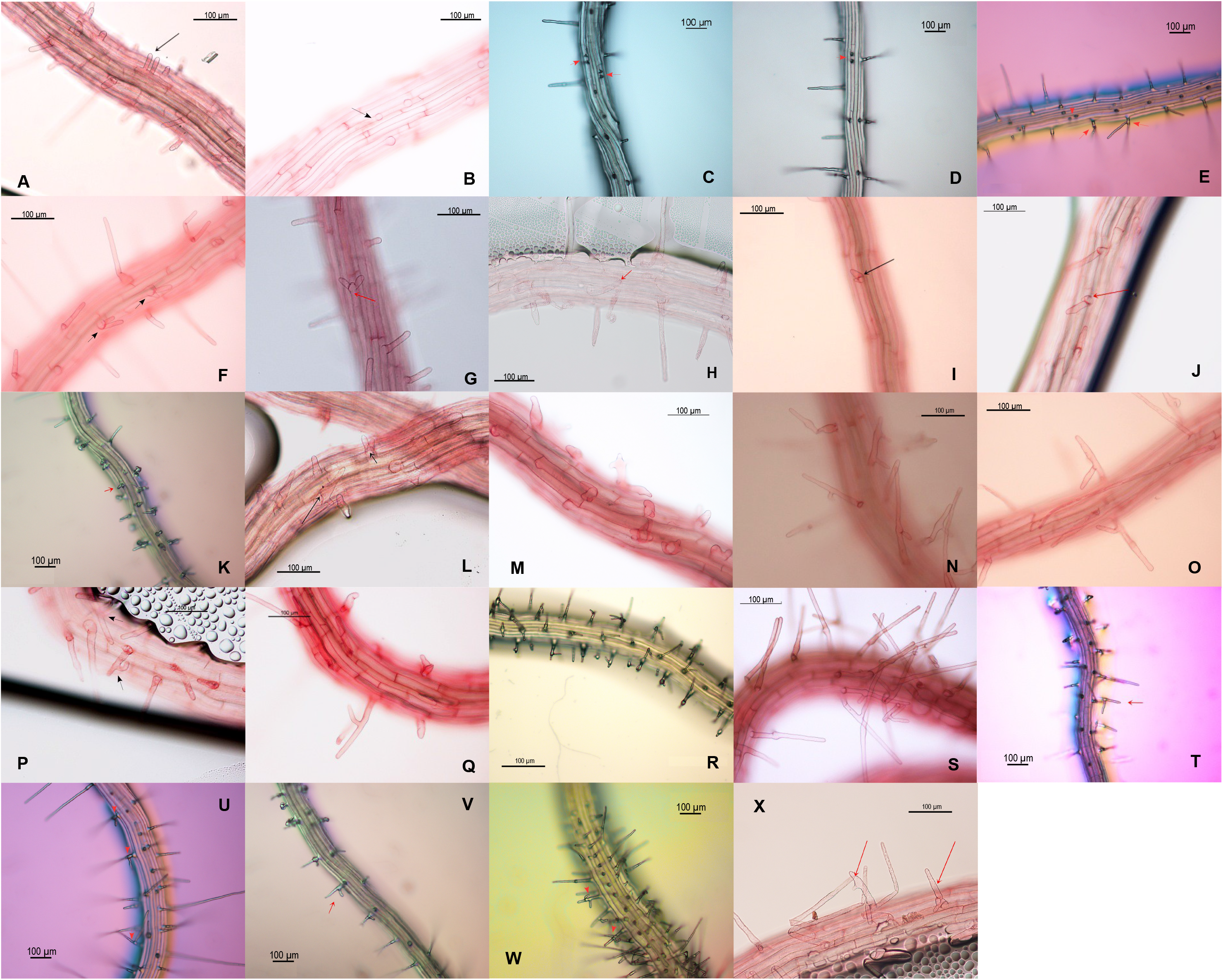
Morphology of trichoblasts and root hairs in root tips of 5-day-old A. thaliana seedlings grown under (or harbouring) excessive levels of GAs/DELLAs. **A)** Two-haired cell in PAC (0.5 μM), 20X; **B)** Cell with two hair bulges in PAC (0.5 μM), 20X; **C)** Two-haired cells in *gai-1,* 10X; **D)** Two-haired cells in *QD,* 10X; Two-haired cells and branched hairs in *UAS::gai-1* x Q2393, 10X; **F)** Two-tipped hairs in PAC (0.5 μM), 20X; **G)** Two-tipped hairs in GA_4_ (1 μM), 20X; **H)** Two-tipped hairs in *gai-1,* 20X; **I)** Two-tipped hairs in *QD,* 20X; **J)** Two-tipped hairs in *pGAI::gai-1:GR* (10 μM DEXA), 20X; **K)** Two-tipped hairs in *UAS::gai-1* x J0121, 10X; **L)** Two-tipped and branched hairs in PAC (0.5 μM), 20X; **M)** Branched hairs in PAC (0.5 μM), 20X; **N)** Branched hairs in *gai-1*, 20X; **O)** Branched hairs in *QD,* 20X; **P)** Branched hairs in *pGAI::gai-1:GR* (10 μM DEXA), 20X; **Q)** Branched hairs in *SCR::gai-1:GR* (MS; leaky line), 20X; **R)** Branched hairs in *UAS::gai-1* x J0951, 20X; **S)** Branched hairs in *UAS::gai-1* x J2812, 20X; **T)** Branched hairs in *UAS::gai-1* x N9142, 10X; **U)** Branched hairs in *UAS::gai-1* x Q2393, 10X; **V)** Branched hairs in *UAS::gai-1* x J0121, 10X; **W)** Branched hairs in *UAS::gai-1* x J0631, 10X; **X)** Branched hairs in *ML1::gai-1*, 20X.

**Fig. 5A.**
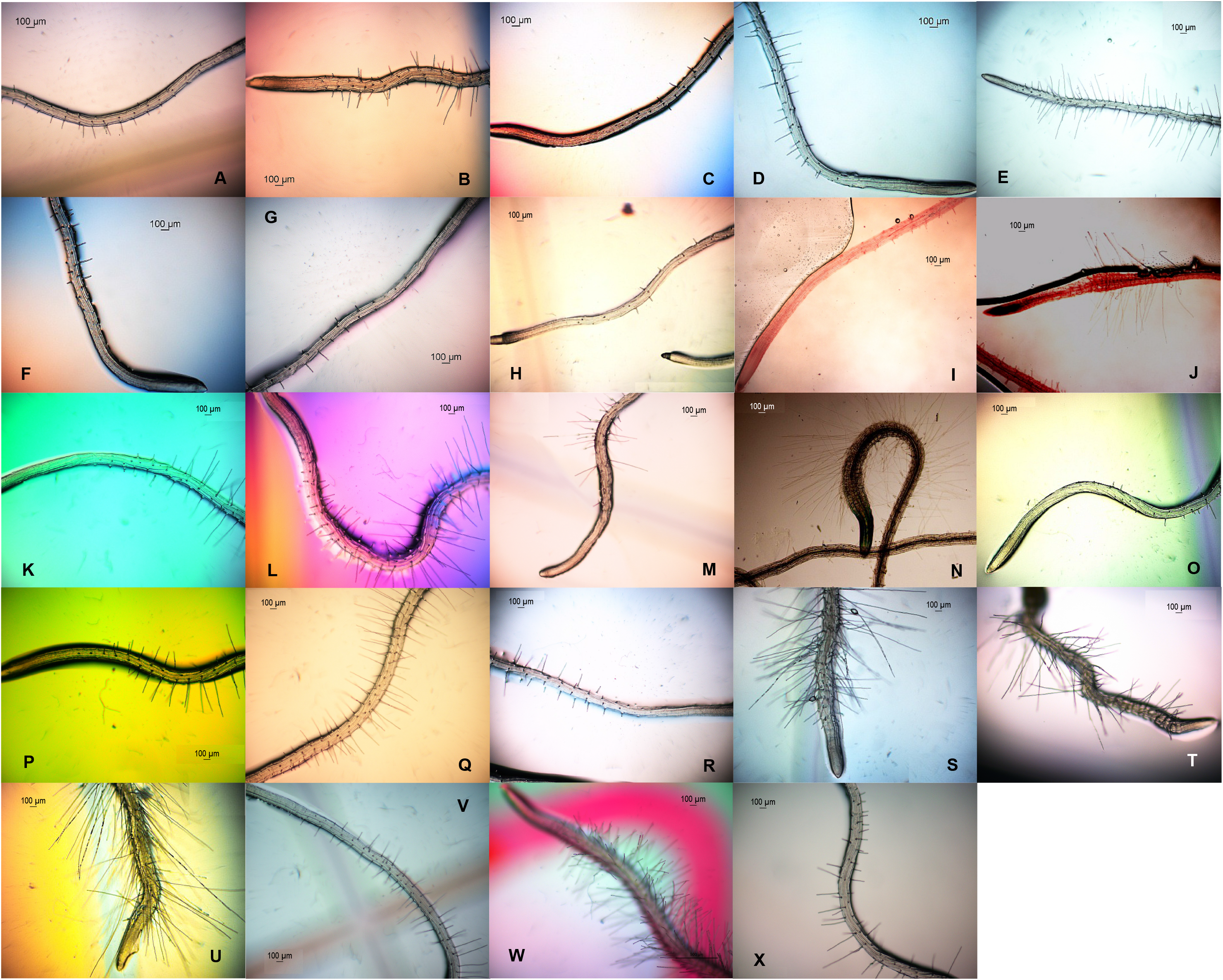
Length and abundance of hairs in root tips of 5-day-old A. thaliana seedlings grown under (or harbouring) excessive levels of GAs/DELLAs. **A)** Col(0) (MS); **B)** Col(0) (0.5 μM PAC); **C)** Col(0) (1 μM GA_4_); **D)** L*er*; **E)** *gai-1*; **F)** *QD*; **G)** *5X*; **H)** GID*1b-ox* (MS; leaky line); **I)** *HSp::gai-1* (24h at 22°C); **J)** *HSp::gai-1* (24h after heat-shock (37°C for 4h)); **K)** *pGAI::gai-1:GR* (30h in MS; leaky line); **L)** *pGAI::gai-1:GR* (30h in 10 μM DEXA); **M)** *SCR::gai-1:GR* (24h in MS; leaky line); **N)** *SCR::gai-1:GR* (24h in 10 μM DEXA); **O)** *UAS::gai-1* x C24; **P)** *UAS::gai-1* x J0951; **Q)** *UAS::gai-1* x J2812; **R)** *UAS::gai-1* x N9142; **S)** *UAS::gai-1* x M0018; **T)** *UAS::gai-1* x J0571; **U)** *UAS::gai-1* x Q2500; **V)** *UAS::gai-1* x"bibr" Q2393; **W)** *UAS::gai-1* x J0631; **X)** *UAS::gai-1* x J0121. Magnification: 4X.

**Fig. 5B.**
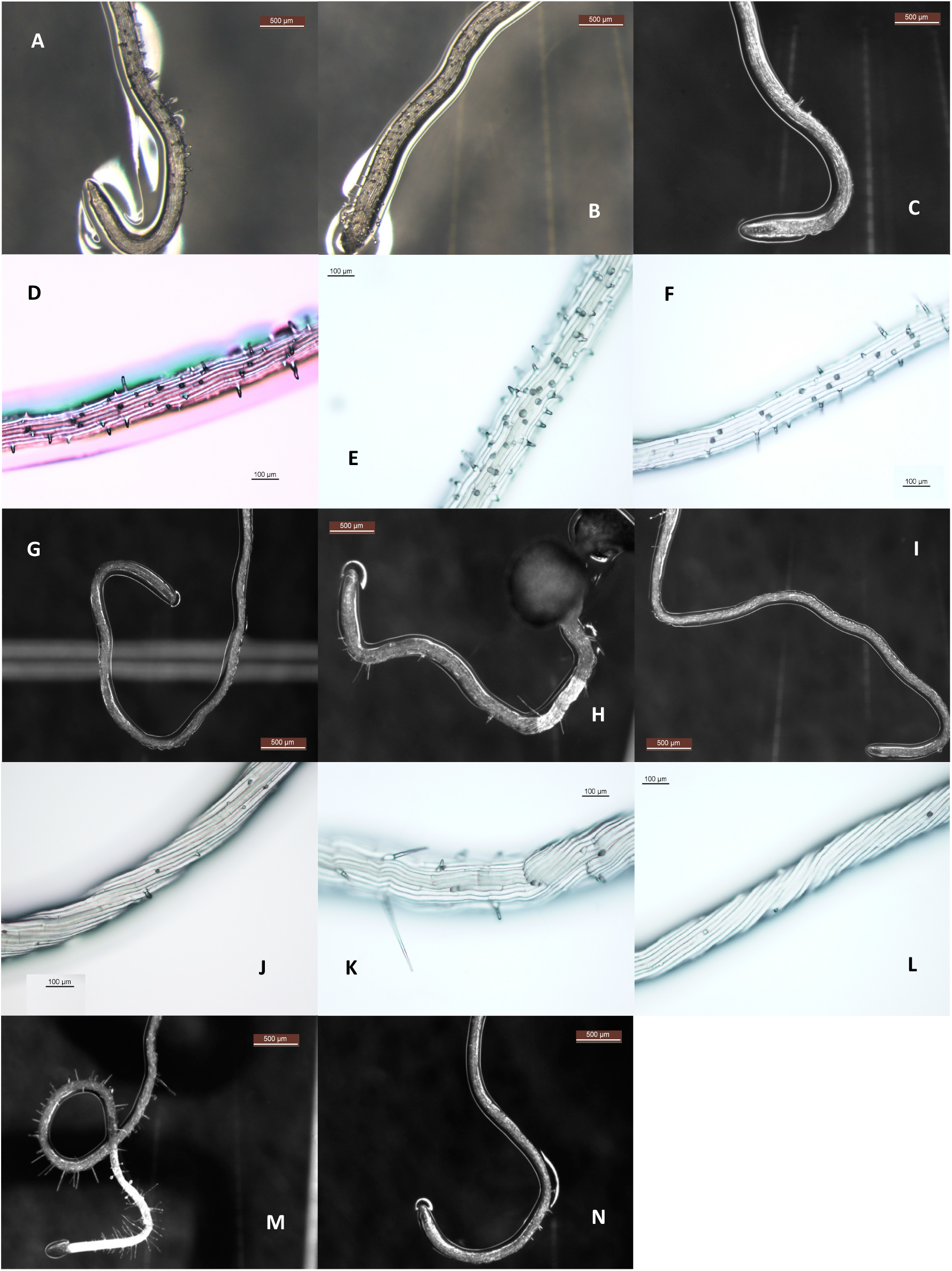
Length and abundance of hairs in root tips of wer, cpc and 35S::CPC mutant seedlings grown for 5 days under excessive levels of GAs/DELLAs. **A)** *wer* mutant (MS), lens; **B)** *wer* mutant (0.5 μM PAC), lens; **C)** *wer* mutant (1 μM GA_4_), lens; **D)** *wer* mutant (MS), microscope; **E)** *wer* mutant (0.5 μM PAC), microscope; **F)** *wer* mutant (1 μM GA_4_), microscope; **G)** *cpc* mutant (MS), lens; **H)** *cpc* mutant (0.5 μM PAC), lens; **I)** *cpc* mutant (1 μM GA_4_), lens; **J)** *cpc* mutant (MS), microscope; **K)** *cpc* mutant (0.5 μM PAC), microscope; **L)** *cpc* mutant (1 μM GA_4_), microscope; **M)** *35S::CPC* mutant (MS), lens; **N)** *35S::CPC* mutant (1 μM GA_4_), lens. Magnification: 2.5 X (lens), 10X (microscope).

**Table 3.** Length and abundance of hairs in root tips of 5-day-old A. thaliana seedlings grown under (or harbouring) excessive levels of GAs/DELLAs. Analyses of hair length and abundance were performed on electron micrographs of root tips of *A. thaliana* seedlings (4X). (*) Seedlings analysed at 24h and 48h after a heat-shock experiment (37°C, 4h). Analyses of hair abundance performed at 31.5X (lens; Control, PAC and GA_4_), 3.2X (lens; L*er*, *gai-1* and *QD*) or 4X (microscope; other mutants and *UAS::gai-1* lines).

|  | Hairs analysed | Root Hair Length (µm) | Roots examined | N° of Root Hairs per field |
| --- | --- | --- | --- | --- |
| Col (0) (MS) | 69 | 209 ± 121 (100%) | 17 | 38 ± 8 (100 %) |
| Col (0) (0.5 µM PAC) | 94 | 270 ± 128 (129 %) | 19 | 54 ± 12 (142 %) |
| Col (0) (1 µM GA<sub>4</sub>) | 37 | 178 ± 93 (85 %) | 18 | 31 ± 8 (82 %) |
| *Ler* | 45 | 201 ± 99 (100 %) | 5 | 43 ± 7 (100 %) |
| *gai-1* | 120 | 397 ± 186 (198 %) | 3 | 56 ± 1 (130 %) |
| *QD* | 25 | 139 ± 85 (69 %) | 3 | 24 ± 5 (56 %) |
| *GID1b-ox* | 14 | 80 ± 25 (40 %) | 6 | 28 ± 8 (88 %) |
| *GID1b-ox* (30 µM GA<sub>3</sub>) | 10 | 64 ± 35 (32 %) | 3 | 18 ± 6 (57 %) |
| *Hsp::gai-1* (22 °C) at 24h (*) | 6 | 55 ± 14 (100 %) | 1 | 18 ± 0 (100 %) |
| *Hsp::gai-1* (37 °C) at 24h (*) | 11 | 405 ± 208 (201 %) | 2 | 32 ± 4 (178 %) |
| *Hsp::gai-1* (37 °C) at 48h (*) | 23 | 411 ± 165 (204 %) | 3 | 83 ± 31 (459 %) |
| *pGAI::gai-1:GR* (MS, 30h) | 40 | 270 ± 118 (100 %) | 4 | 56 ± 7 (100 %) |
| *pGAI::gai-1:GR* (0.5 µM DEXA, 30h) | 57 | 314 ± 177 (116 %) | 8 | 79 ± 17 (142 %) |
| *SCR::gai-1:GR* (MS, 3d) | 30 | 245 ± 87 (100 %) | 3 | 49 ± 20 (100 %) |
| *SCR::gai-1:GR* (0.5 µM DEXA, 3d) | 35 | 507 ± 173 (207 %) | 5 | 76 ± 31 (154 %) |
| *wer* (MS) | 24 | 192 ± 88 (100 %) | 3 | 91 ± 12 (100 %) |
| *wer* (0.5 µM PAC) | 8 | 243 ± 134 (127 %) | 3 | 125 ± 29 (137 %) |
| *wer* (1 µM GA<sub>4</sub>) | 6 | 133 ± 23 (70 %) | 3 | 70 ± 5 (77 %) |
| *cpc* (MS) | 7 | 104 ± 29 (100%) | 3 | 17 ± 1 (100 %) |
| *cpc* (0.5 µM PAC) | 9 | 213 ± 92 (204 %) | 3 | 18 ± 2 (106 %) |
| *cpc* (1 µM GA<sub>4</sub>) | 8 | 88 ± 51 (85 %) | 7 | 11 ± 3 (65 %) |
| *UAS::gai-1* x C24 (control) | 20 | 161 ± 105 (100 %) | 2 | 48 ± 23 (100 %) |
| *UAS::gai-1* x J0951 | 34 | 240 ± 118 (149 %) | 3 | 49 ± 13 (101 %) |
| *UAS::gai-1* x J2812 | 59 | 243 ± 118 (151 %) | 9 | 77 ± 26 (161 %) |
| *UAS::gai-1* x J0571 | 25 | 586 ± 273 (364 %) | 2 | 60 ± 11 (125 %) |
| *UAS::gai-1* x M0018 | 90 | 685 ± 195 (425 %) | 10 | 92 ± 26 (192 %) |
| *UAS::gai-1* x Q2500 | 37 | 680 ± 189 (422 %) | 2 | 96 ± 12 (200 %) |
| *UAS::gai-1* x Q2393 | 48 | 272 ± 146 (169 %) | 4 | 67 ± 26 (140 %) |
| *UAS::gai-1* x N9142 | 21 | 195 ± 97 (121 %) | 2 | 30 ± 1 (63 %) |
| *UAS::gai-1* x J0121 | 47 | 233 ± 120 (145 %) | 5 | 67 ± 13 (140 %) |
| *UAS::gai-1* x J0631 | 8 | 386 ± 129 (240 %) | 2 | 96 ± 15 (200 %) |

High levels of GAs/DELLAs also altered the abundance of hairs in the radial dimension of the root tips (Tables 4 and 5). The number of root hairs per root cross section, calculated as the summary of root hairs at the trichoblast and atrichoblast positions (or the summary of root hairs and ectopic root hairs per root cross section) increased under excessive DELLAs (PAC, *gai-1*) but decreased in the *5X* mutant (Table 5). On the other hand, the number of root non-hairs per root cross section, calculated as the summary of root non-hairs at the atrichoblast and trichoblast positions (or the summary of root non-hairs and ectopic root non-hairs per root cross section), decreased under excessive DELLAs, but experienced an enhancement in the *5X* mutant (Table 5). Thus, the estimated abundance of root hairs in the radial dimension of root tips seemed to increase under excessive DELLAs, but to decrease under excessive GAs.

**Table 4.**
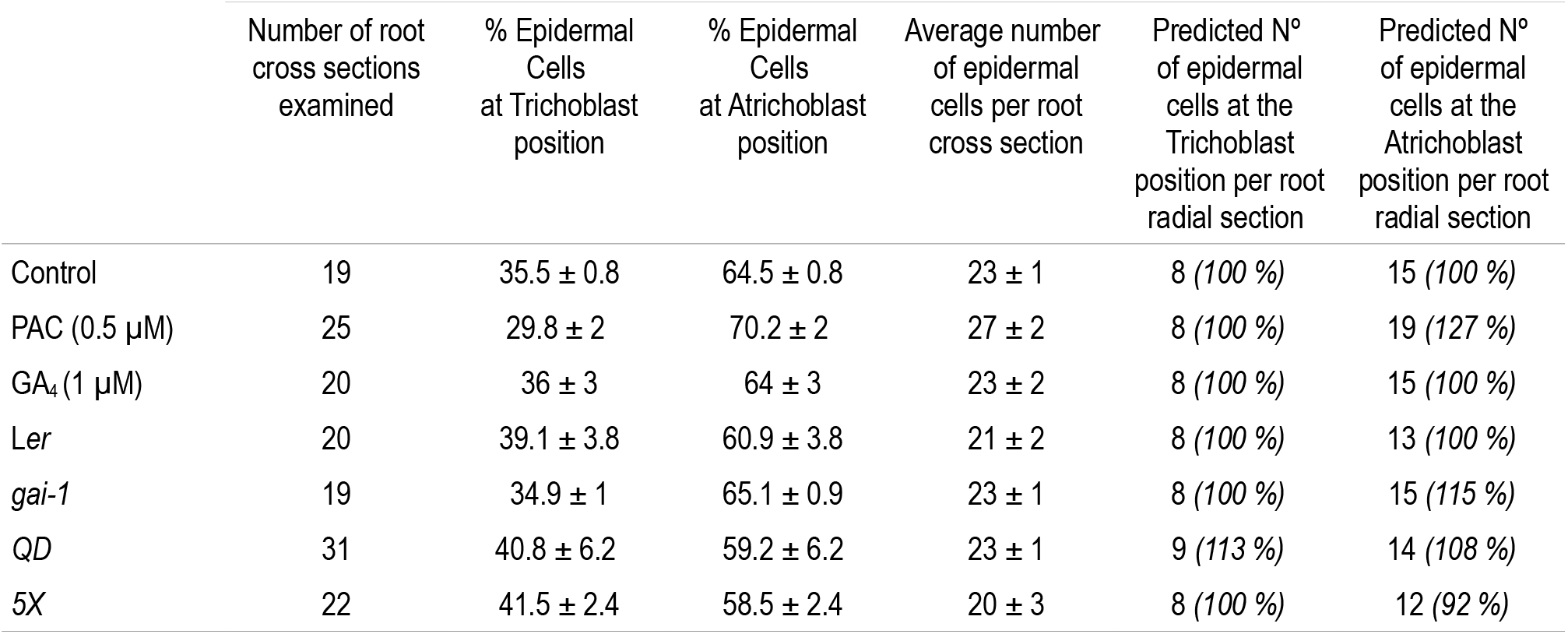
Percentage and estimated number of epidermal cells at the trichoblast/atrichoblast positions per root cross section in 5-day-old A. thaliana seedlings grown under (or harbouring) excessive levels of GAs/DELLAs. Analyses performed on electron micrographs of cross sections of resin-embedded roots (40X).

**Table 5.** Estimated number of root hairs and non-hairs per root cross section in root tips of 5-day-old A. thaliana seedlings grown under (or harbouring) excessive levels of GAs/DELLAs. Calculations were made by considering the data of Table 1 (distribution of hair and non-hair cells at the trichoblast/atrichoblast positions), Table 2 (percentage of ectopic root hair/non-hair cells at the atrichoblast/trichoblast positions) and Table 4 (average number of epidermal cells per root cross section, estimated number of epidermal cells at the trichoblast position per root cross section, and estimated number of epidermal cells at the atrichoblast position per root cross section). Estimated number of root hairs per root cross section = [hairs at the trichoblast position + hairs at the atrichoblast position]. Estimated number of root non-hairs per root cross section = [non-hairs at the atrichoblast position + non-hairs at the trichoblast position].

|  | Trichoblast position |  | Atrichoblast position |  | Estimated N° of Root Hairs per root cross section | Estimated N° of Non-root hairs per root cross section |
| --- | --- | --- | --- | --- | --- | --- |
|  | Hairs per root cross section | Non-hairs per root cross section | Hairs per root cross section | Non-hairs per root cross section |  |  |
| Control | 8 | 0 | 0 | 15 | 8 (100 %) | 15 (100 %) |
| PAC (0.5 µM) | 6 | 2 | 7 | 12 | 13 (163 %) | 14 (93 %) |
| GA<sub>4</sub> (1 µM) | 6 | 2 | 2 | 13 | 8 (100 %) | 15 (100 %) |
| Ler | 8 | 0 | 1 | 12 | 9 (100 %) | 12 (100 %) |
| *gai-1* | 7 | 1 | 6 | 9 | 13 (144 %) | 10 (83 %) |
| QD | 7 | 2 | 3 | 11 | 10 (111 %) | 13 (108 %) |
| 5X | 5 | 3 | 0 | 12 | 5 (56 %) | 15 (125 %) |

## 4. DISCUSSION

### 4.1. The GAs/DELLAs might regulate the root hair patterning in seedlings of *A. thaliana*

Whereas the role of GAs/DELLAs in the production and distribution of leaf hairs has been well studied (Telfer *et al.,* 1997; Perazza *et al.,* 1998), their hypothetical function in the determination and arrangement of root hairs has not been examined up to date. To this aim, the effects of high levels of GAs/DELLAs on the spatial gene expression of the hair (CPC) and non-hair (GL2, WER and EGL3) markers of root epidermal cell fate, as well as on the distribution of root hairs, were analysed in seedlings of *A. thaliana*. Results showed that excessive levels of GAs/DELLAs impaired the spatial gene expression of the root hair/non-hair epidermal cell fate markers and disarranged the normal distribution of root hairs, what suggested that the GAs/DELLAs might be involved in regulating the root hair patterning in seedlings of *A. thaliana.* In fact, stable or inducible mutants with low (*gai-1, HSp::gai-1*, *pGAI::gai-1:GR*, *SCR::gai-1:GR*) or high (*QD*, *5X*, *GID1b-ox*) levels of GAs showed not only a random expression of *GL2* at the MZ and EZ of the root, known as the cell fate-decision zones (Pernas *et al.,* 2010), but also a disarrangement of the root hairs. Because neither the *GL2* spatial expression nor the distribution of root hairs suffered changes when the gai-1 DELLA was over-expressed at the root epidermis (*ML1::gai-1* x *GL2pro::GUS*, *ML1::gai-1* and *UAS::gai-1* x J0951 transgenic lines), it was concluded that the GAs/DELLAs do not seem to affect the root hair patterning in *A. thaliana* seedlings by acting on this cell layer of the root, but on tissues placed underneath. In fact, expression of *gai-1* at the cortex, endodermis or pericycle of the root MZ altered the root hair patterning.

Interestingly, expressing *CPC* at the stele rescues the phenotype of the hairless *cpc* mutant, what suggests that epidermal cell differentiation might be controlled from the internal tissues of the root (Rishmawi *et al.,* 2014). Therefore, the results of this study suggest that, as it was previously reported for auxins, ET, ABA, NO, BRs and SLs (Schiefelbein, 2003), the GAs/DELLAs might regulate the root hair patterning in seedlings of *A. thaliana* independently from the gene network for the root epidermal cell fate specification, although confirmatory studies might be required.

The reason why excessive levels of GAs/DELLAs disarranged the root hair patterning in *A. thaliana* seedlings might have been, in part, related to their effects on the cytoskeleton of MT. The MT cytoskeleton, consisting in polymers of α and β tubulin, is essential for the appropriate distribution of positional signals during development (Schiefelbein, 2003). Also, the orientation of MT participates in the determination of epidermal cell fate (Pietra, 2014). Thus, MT lay randomly in trichoblasts but transversally in atrichoblasts (Dugardeyn and Van der Straeten, 2008). Hormone-induced reorganization of MT is also necessary for root hair initiation (Bao *et al.,* 2001; Schiefelbein, 2003). Interestingly, the GAs/DELLAs regulate MT organization by interacting with prefoldin, a protein required for the folding of tubulin (Locascio *et al.,* 2013). As a result of this interaction, MT are organized in the presence of GAs, like in root or mesocotyl epidermal cells, and disorganized in the presence of DELLAs (Perazza *et al.,* 1998; Bouquin *et al.,* 2002; Locascio *et al.,* 2013). On the other hand, mutants impaired in MT assembly have an altered root hair patterning (Bouquin *et al.,* 2002). The *lue1* mutant, which lacks a MT-severing and cell wall (CW) biosynthesis-related katanin protein, and whose MT are disorganized, is allelic to ectopic root hair 1 ( *erh1*) and has an altered root hair patterning (Bouquin *et al.,* 2002; Webb *et al.,* 2002). In addition, *lue1* presents an inappropriate regulation of the GA biosynthesis-related AtGA*20ox* activity and responds to GAs (Schneider *et al.,* 1997; Bouquin *et al.,* 2002).

Ectopic root hairs have also been described in TUA6/AS transgenic lines under-expressing α-tubulin genes, in plants treated with MT polymerization-inhibiting drugs or with trichostatin A (TSA, a histone deacetylase (HDA) inhibitor), during the inducible expression of MT-interacting phospholipase-D (PLD) activity, as well as in mutants of MT severing/reorganization-related proteins, such as HDA, COBRA, SABRE and katanin p60 (Schiefelbein *et al.,* 1997; Bao *et al.,* 2001; Bouquin *et al.,* 2002; Sedbrook, 2004; Wang, 2005; Xu *et al.,* 2005; Li *et al.,* 2006, 2015; Chen *et al.,* 2015; Pietra *et al.,* 2015). In fact, the katanin complex is required for the specification of root epidermal cells (Webb *et al.,* 2002). In addition, the katanin P60-related alteration of MT organisation affects the composition and deposition of the CW (Sedbrook, 2004). Histone deacetylation also participates in cellular patterning, because TSA-induced histone acetylation modifies GL2, WER and CPC expression and localization and induces ectopic root hairs (Xu *et al.,* 2005; Cui and Benfey, 2009). Lack of SABRE function equally destabilizes the expression of cell fate markers, including WER and GL2 (Pietra *et al.,* 2015). In addition, a delocalized expression of GL2 has been documented for the *jkd* (jackdaw) and *scm* (scrambled) mutants (Hassan *et al.,* 2010; Pietra, 2014).

Therefore, the MT participate in cell identity specification (Webb *et al.,* 2002). Cell identity, in turn, mediates the root responses to abiotic stress (Dinneny *et al.,* 2008). Thus, ectopic root hairs and non-hairs have been described in *A. thaliana* seedlings exposed to gamma irradiation, Cd or As, and during P deficiency, although without quantitative changes in the *WER* and *GL2* expression (Ma *et al.,* 2001; Nagata *et al.,* 2004; Yang *et al.,* 2007; Bahmani *et al.,* 2016). Moreover, stress down-regulates the actin and tubulin gene expression (Sánchez-Calderon *et al.,* 2013). In turn, a reduced expression of the α-tubulin gene results in MT disassembly, with MT laying in an aberrant way, and in their reorganization (Bao *et al.,* 2001).

Consequently, the root hair patterning responds to environmental signals (Salazar-Henao *et al.,* 2016). For instance, the photoperiod and thermoperiod control the root hair patterning in tomato (Tsai *et al.,* 2004). Interestingly, the GAs participate in thermotolerance (Alonso-Ramirez *et al.,* 2009). Thus, the results of this study suggest that the GAs/DELLAs might regulate, in part, the root hair patterning in *A. thaliana* seedlings by altering MT organization. In root cells, excessive levels of DELLAs might disorganize the cytoskeleton of MT, thereby impairing the link between positional signals and cell fate, whereas excessive levels of GAs might stabilize it.

Results of this study also point at a possible role for the DELLAs in regulating the root hair patterning in response to nutritional deficiencies. The random disposition of root hairs under excessive levels of DELLAs might favour the foraging of scarce or non-mobile minerals in deficient soils. Thus, altering the root hair patterning by modulating the levels of GAs/DELLAs might constitute a mechanism used by plants for increasing the possibilities of acquiring non-available minerals, such as P or Fe, in deficient soils. In fact, plant deficiencies in P, B or Fe disarrange the root hair patterning and induce ectopic root hairs (Schmidt *et al.,* 2000; Péret *et al.,* 2011; Janes *et al.,* 2018). Moreover, low availability of P increases the levels of DELLAs and reduces the levels of GAs in roots (Jiang *et al.,* 2007).

Results of this study also suggest that the GAs/DELLAs might affect the root hair patterning in *A. thaliana* seedlings by acting not at the epidermis, where the gene network for the root hair/non-hair epidermal cell fate operates, but at tissues placed underneath (cortex, endodermis and pericycle). However, confirmatory studies are still needed to uncover why the epidermal expression of *gai-1* did not modify the root hair patterning in *A. thaliana* seedlings, in spite that the DELLAs promote the disorganization of MT in root epidermal cells. Moreover, the fact that only one DELLA (gai-1) was over-expressed in this study, and that expression of *gai-1* at the epidermis (*ML1::gai-1*, J0951 ≫ *gai-1*) induced longer and branched root hairs, suggests that the effects of the GAs/DELLAs on the root epidermal cells and/or the root hair patterning in seedlings of *A. thaliana* might be different depending on the particular concentration at which these hormones might be present.

### 4.2. The GAs/DELLAs might regulate the shape, length and abundance of hairs in root tips of *A. thaliana* seedlings

Supra-physiological levels of GAs/DELLAs in roots of *A. thaliana* seedlings also induced two-haired cells, two-tipped hairs and branched hairs. Multiple hairs per root epidermal cell, two-tipped hairs and branched hairs have also been reported in the SUPERCENTIPEDE (*scn1*) mutant, with supernumerary root hair initiation sites, in TUA6/AS *A. thaliana* transgenic lines under-expressing α-tubulin genes, in root hair defective 3, 4 and 6 (*rhd3*, *rhd4*, *rhd6*) and *PLD* mutants, in plants treated with MT-depolymerizing oryzalin, MT-disorganizing 1-butanol (a PLD-inhibitor) or MT-stabilizing taxol, in ROP2 (proteins controlling MT organization) over-expressing plants, and in plants subjected to Fe or NO_3^−^_ deficiency (Schiefelbein and Somerville, 1990; Schiefelbein *et al.,* 1993; Masucci and Schiefelbein, 1994; Gilroy and Jones, 2000; Schmidt *et al.,* 2000; Bao *et al.,* 2001; Foreman and Dolan, 2001; Grierson and Schiefelbein, 2002; Jones *et al.,* 2002; Gardiner *et al.,* 2003; Müller and Schmidt, 2004; Carol and Dolan, 2006; Ishida *et al.,* 2008; Shin *et al.,* 2011; Pietra, 2014). Interestingly, hormone-induced reorganization of MT is required for the morphogenesis of root hairs (Bao *et al.,* 2001; Schiefelbein, 2003). In turn, the phenotype of root hair branching, due to changes in actin distribution and dynamics, has been related to the induction of genes for GA biosynthesis and CW modification, and reported during legume-rhizobium symbiosis (i.e., soybean infected with *Bradyrhyzobium japonicum*), in plants treated with MT-inhibiting drugs, and in mutants of genes necessary for a correct growth of root hairs, such as *TIP1* (involved in the biosynthesis of CW components and probably in the arrangement of actin filaments) and *RHD3* (Schiefelbein and Somerville, 1990; Schiefelbein *et al.,* 1993; Bao *et al.,* 2001; Salazar-Henao *et al.,* 2016).

The disruption of MT also affects trichome branching (Gilroy and Jones, 2000), as actin regulates the shape and growth of trichomes (Rodríguez-Serrano *et al.,* 2014). In addition, the GAs promote trichome branching and influence CW growth (Telfer *et al.,* 1997; Perazza *et al.,* 1998). Thus, the *spy5* mutant (with high levels of GAs and which also displays ectopic root hairs) has over-branched trichomes (Perazza *et al.,* 1998; Mutanwad *et al.,* 2020). On the other hand, during trichome development, the number of branches and the level of endo-reduplication, which is induced by GAs, are closely related (Perazza *et al.*, 1998; Kondorosi *et al.,* 2001).

In this study, excessive levels of DELLAs in *A. thaliana* seedlings also induced longer root hairs near the root tip. Interestingly, nutrient availability prevents root hair elongation (Tsai *et al.,* 2004), whereas deficiencies in P, B or Mg induce root hair elongation, being the higher levels of DELLAs the mediators of the extra-elongation of root hairs (Péret *et al.,* 2011; Liu *et al.*, 2018). Elongated root hairs have also been described in plants exposed to gamma irradiation, Cd or As, as well as in polyploids (Nagata *et al.,* 2004; Setter *et al.,* 2015; Bahmani *et al.*, 2016; Salazar-Henao *et al.,* 2016). Conversely, shorter root hairs have been reported in mutants of the *TIP1*, *PLDξ1-PLDξ2*, and *RSL4* (a component of GAs signalling) genes (Schiefelbein *et al.,* 1993; Li *et al.,* 2006; Péret *et al.,* 2011). Moreover, the GAs are necessary for root hair elongation, as the *ga* 1-3 mutant (deficient in GAs) produces shorter root hairs (Péret *et al.,* 2011). However, the GAs might act at a later stage of root hair development, as apparently, in this study, high levels of GAs did not stimulate root hair elongation near the root tip as much as the high levels of DELLAs did. Therefore, the changes induced, in this study, by excessive levels of GAs/DELLAs on the shape and length of root hairs in seedlings of *A. thaliana* might have been related to the effect of these hormones on the MT cytoskeleton and/or the CW biosynthesis of the root epidermal cells.

Regarding root hair abundance, it is known that nutrient availability inhibits root hair production (Tsai *et al.,* 2004). Excess of Na^+^ reduces root hair abundance (Dinneny *et al.,* 2008), whereas deficiencies in P, Fe or B increase the frequency of root hairs, mainly by inducing ectopic root hair cells (Schiefelbein, 2003; Martín-Rejano *et al.,* 2011; Péret *et al.,* 2011; Shin *et al.,* 2011; Salazar-Henao *et al.,* 2016; Janes *et al.,* 2018). An increased density of root hairs has also been reported in ROP2 over-expressing plants, in *arm (ctl1*; cellulose biosynthesis-related) and *sabre* mutants, in plants exposed to Cd, Vd or As, and in polyploids (Jones *et al.*, 2002; Pietra, 2014; Lin *et al.,* 2015; Bahmani *et al.*, 2016; Salazar-Henao *et al.,* 2016). Interestingly, the levels of GAs determine trichome number (Perazza *et al.,* 1998). In turn, HDA19 controls the response of the root hair density to low P (Chen *et al.,* 2015).

Because of the GAs/DELLAs are involved in plant stress responses (Alonso-Ramírez *et al.,* 2009), the results of this study suggest that these hormones might have a role in regulating the response of the root hair abundance to nutrient availability. In fact, in this study, root hairs near the root tip were denser and longer under excessive DELLAs, but scarcer and shorter under excessive GAs. With this respect, it is known that root hairs grow closer to the root MZ under mechanic stress or B deficiency (Okamoto *et al.,* 2008; Martín-Rejano *et al.,* 2011). Also, the abundance and length of root hairs respond to environmental signals (Salazar-Henao *et al.,* 2016). Light signalling, for instance, influences root hair length (Grierson and Schiefelbein, 2002). In turn, the photo-period conditions affect the biosynthesis and/or sensibility of GAs (Telfer *et al.,* 1997).

As PLD inhibitors break the organization of MT, which is essential for the correct directionality, elongation and morphology of root hairs (Gardiner *et al.,* 2003), then, the morphological alterations of root hairs observed in this study point to a possible impairment, by excessive levels of GAs/DELLAs, of the actin microfilaments, the cytoskeleton of MT, and the ROP GTPase proteins. In fact, hair cell morphogenesis requires α-tubulin and Rho-like GTPase activity, which, in turn, interacts with katanin P60 to promote MT ordering (Foreman and Dolan, 2001; Lin *et al.,* 2013). Moreover, the SABRE protein (involved in MT organisation and the stabilization of epidermal patterning factors) acts upstream of ROPs (Pietra, 2014).

## ACKNOWLEDGEMENTS

Special thanks to Dr. Miguel Ángel Blázquez and Dr. David Alabadí for supporting the writing of this paper. This study was performed at the Blázquez-Alabadí laboratory (Hormone Signalling and Plant Plasticity Group) of the Instituto de Biología Celular y Molecular de Plantas (IBMCP)-UPV-CSIC, Valencia, Spain. Thanks also to Dr. Benedicte Desvoyes for the training in the hair/non-hair cell counting (Dr. Crisanto Gutiérrez Lab). I. McCarthy-Suárez also acknowledges a JAE-Doc postdoctoral fellowship ((2008-2011), C.S.I.C., Valencia, Spain). This paper is dedicated to the loving memory of Dr. Francisco Culiáñez-Macià, Principal Investigator at the IBMCP-UPV-CSIC, Valencia, Spain.

## Notes

### Competing Interest Statement

The authors have declared no competing interest.

